# Persistence length of short homopolymeric single-stranded DNA sequences in polyvalent cations

**DOI:** 10.64898/2026.08.02.742318

**Authors:** Balaka Mondal, D. Thirumalai

## Abstract

We calculated the persistence length (*l*_p_) of short single-stranded DNA (ssDNA) homopolymers in polyvalent cations using simulations based on the sequence-dependent Three Interaction Site (TIS) model. The model accounts for base stacking, and electrostatic interactions that are treated using the Coulomb potential with explicit ions. In Mg^2+^ and Ca^2+^ cations, *l*_p_ of dT_30_ (T is thymine) varies as 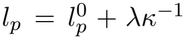, where 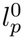 is the bare persistence length, *λ* is a constant, and *κ* is the inverse Debye length. The *κ*^−1^ dependence is surprising because it was derived for long flexible polyelectrolytes in which the charges interact via the Debye-Hückel potential. Strong stacking between the bases dA_30_ (A is adenine) increases the bare persistence length of dA_30_ (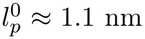 and ≈ 0.4 nm for dT_30_), which is comparable to the electrostatic persistence length. Consequently, neither the flexible (*κ*^−1^) nor the stiff (*κ*^−2^) limits accounts for the simulation results for dA_30_. Interestingly, *l*_p_ for both the sequences is nearly independent of the tetravalent spermine (Spm^4+^) concentration above 0.1 mM. The number of bound Spm^4+^ increases only gradually, a behavior not captured by simple binding models. A qualitative picture, based on the restricted access to the phosphate groups due to excluded volume of the anisotropic Spm^4+^, rationalizes these results. The predicted dependence of *l*_p_ on spermine concentration awaits experimental test.

## INTRODUCTION

Biological molecules, such as DNA and RNA, are highly charged. In general, including polyelectrolytes (referred to as PE), the persistence length (*l*_*p*_) is a measure of their sizes. However, the dependence of *l*_*p*_ on the ionic concentration is difficult to calculate because of the interplay of several length scales [1, 2]. If the PE is intrinsically stiff and if the interaction between the charges is given by the Debye-Hückel potential, then the dependence of *l*_*p*_ is well described by the Odijk-Skolnick-Fixman (OSF) theory [3, 4]. The OSF theory shows that 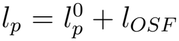, where 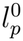 is the bare persistence length. The electrostatic contribution, *l*_*OSF*_ , is given by,

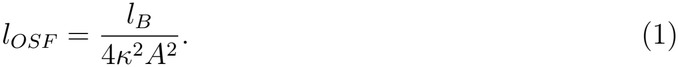

In Eq. 1, the Bjerrum length, 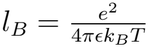 (*e* is the unit of charge, *ϵ* is the dielectric constant, *k*_*B*_ is the Boltzmann constant, and *T* is the temperature), *A* is the distance between the charges on the PE, and *κ* is the inverse screening length (*κ*^2^ = 4*πl*_*B*_*I* where *I* is the ionic strength).

It is known that Eq. 1 is valid when *A*, the distance between the charges on the PE is small 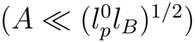, which is satisfied in double stranded DNA for instance. The situation is more complicated if the PE is intrinsically flexible 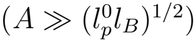, as is the case in single stranded DNA (ssDNA) and RNA [5, 6]. Variational theories [7–10] have established that for flexible PEs, the electrostatic persistence length, *l*_*e*_ scales as,

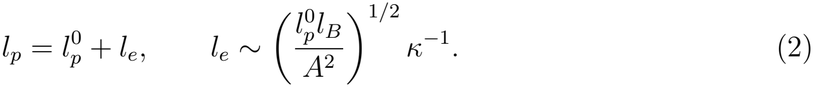

Note that Eq. 1 predicts that *l*_*OSF*_ *∼ I*^−1^, whereas Eq. 2 predicts that *l*_*e*_ *∼ I*^−1*/*2^. The *κ*^−1^ dependence is predicted when the electrostatic contribution exceeds the bare persistence length. The OSF result is recovered in the opposite limit [11]. When the two are comparable, it is difficult to unambiguously determine dependence of *l*_*p*_ on the cation concentration.

The assumption that Debye-Hückel interactions between charged species, which is made in all the theories quoted above, is unlikely to be valid for divalent cations [12] and even less so for spermine, a tetravalent cation at neutral pH. In addition, for ssDNA, we expect that there ought to be differences between sequences, such as polyT and polyA because of differences in the stacking propensity. To assess the applicability of the PE theories and to investigate sequence effects, we simulated ssDNA sequences dT_30_ and dA_30_ in Mg^2+^ and Ca^2+^ as well as in Spm^4+^ in the presence of 20 mM NaCl. We make two unexpected predictions. (1) The persistence length of the sequences varies as 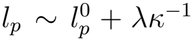 (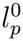 and the dimensionless constant *λ* are dependent on sequence and 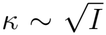 where *I* is the ionic strength). This result is surprising because the variation, *l*_*p*_ *∼ κ*^−1^, was derived for long flexible PE in which the electrostatic interactions were treated using the DH interaction, which are deemed to be inadequate for divalent cations [12, 13]. Because of strong base stacking, the bare persistence length of dA_30_ is comparable to the electrostatic persistence length. The value of 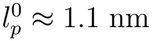 in dA_30_ and *l*_*OSF*_ ≈ 0.5–2.0 nm, so 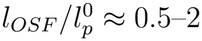. Thus, dA_30_ maybe in the crossover regime, where neither limiting form applies quantitatively. (2) Strikingly, *l*_*p*_ is almost independent of Spm^4+^ concentration above 0.1 mM for both the sequences, which qualitatively differs from the behavior in divalent cations. There is no theory for this observation although simulations suggest that, due to the size of spermine, only a fixed number of Spm^4+^ cations bind to the phosphate groups at all concentrations. The prediction for *l*_*p*_ on Spm^4+^ concentration awaits future experiments.

## METHODS

### Three-Interaction-Site (TIS) model with explicit ions

We used the three-interaction-site (TIS) coarse-grained model for nucleic acids [13–16] in which monovalent Na^+^, divalent (Mg^2+^ and Ca^2+^) and tetravalent cations Spm^4+^ are treated explicitly (see Figure 1). In the TIS representation, each nucleotide is represented by three spherical beads placed at the center of mass of phosphate P, sugar S and base B. Ions are modeled as single spherical beads. The energy function for homopolymeric ssDNA is given by,

**FIG. 1:**
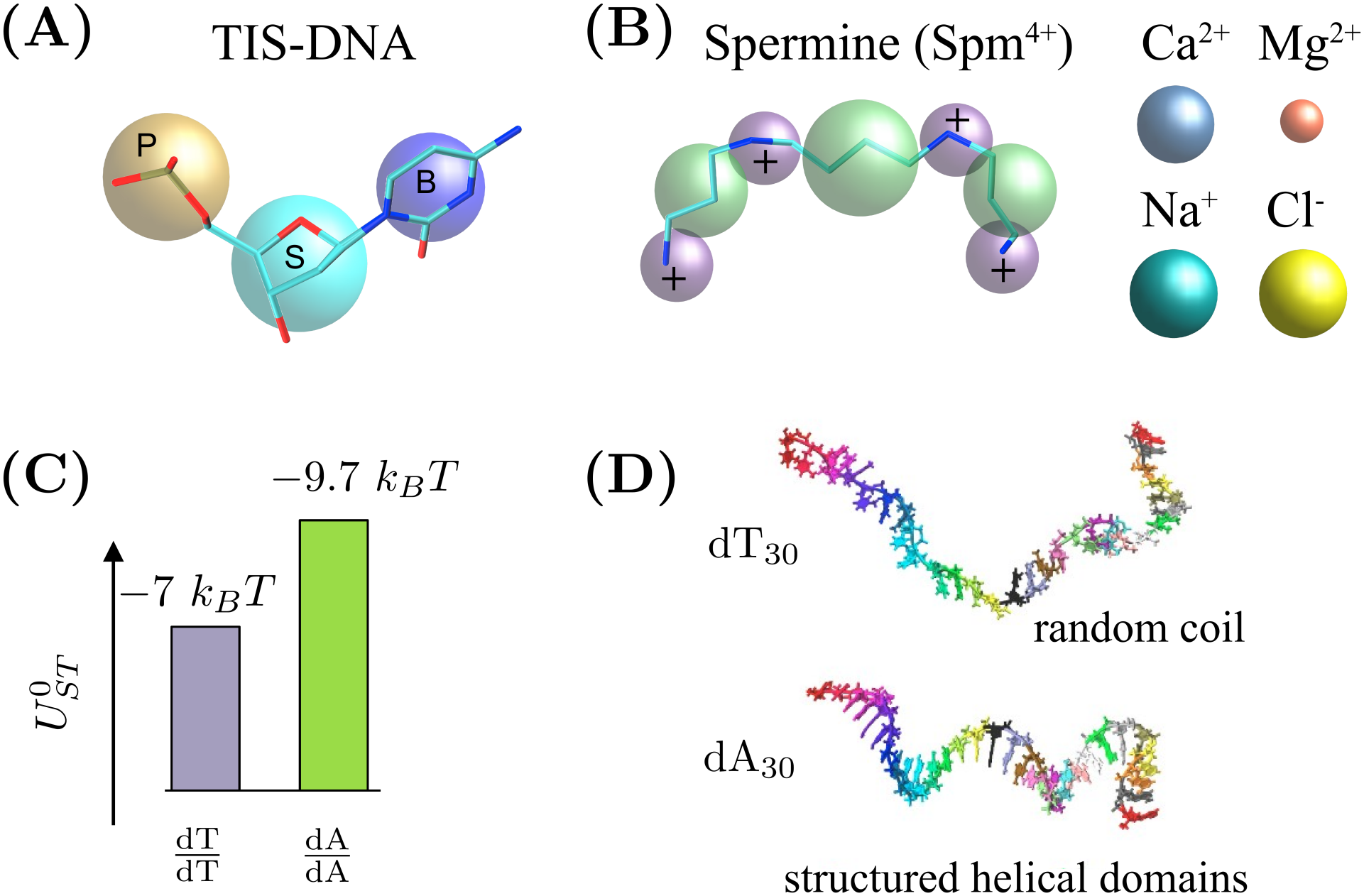
Three-interaction-site (TIS) model for DNA. (A) Each nucleotide is represented by three beads, placed at the center of sugar (cyan), phosphate (tan) and base (blue) groups. (B) Monovalent (Na^+^, Cl^−^) and divalent (Mg^2+^, Ca^2+^) ions are modeled as single coarse-grained beads. The tetravalent Spm^4+^ is represented using seven coarse-grained beads. Purple beads represent charged amine groups and green beads correspond to alkyl groups. (C) Base stacking energies 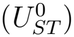 between thymine (dT) and adenine (dA) bases. (D) Random coil-like structure for dT_30_ and a helical conformation for dA_30_, reflecting the differences in the strength of the stacking interactions.

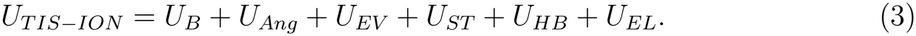

The potentials for bond stretch (*U*_*B*_) and bond angle (*U*_*Ang*_) are given by, *U*_*B*_ = *k*_*b*_(*r* − *r*_0_)^2^ and *U*_*Ang*_ = *k*_*θ*_(*θ* − *θ*_0_)^2^, where *r*_0_ and *θ*_0_ are the equilibrium bond length and bond angle, respectively, which were taken from an ideal B-form DNA helix. The values of *k*_*b*_, *k*_*θ*_, *r*_0_, and *θ*_0_, are listed elsewhere [15]. Excluded volume repulsion between sites *i* and *j* (DNA sites or ions) separated by a distance, *r* (in nm), is modeled as,

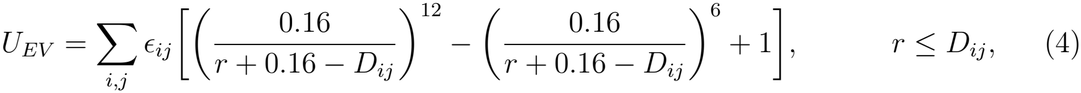

where *D*_*ij*_ = *R*_*i*_ + *R*_*j*_ and 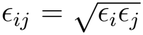. The values of radius *R* and *ϵ* for the ions and DNA are listed in Table S1 of the Supplementary Information (SI). When both the interacting sites belong to DNA, *D*_*ij*_ is taken to be 0.32 nm [17]. Ions are modeled as charged spheres in implicit solvent, interacting via Coulomb and short-range excluded-volume potentials. We use 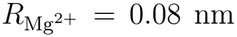 and 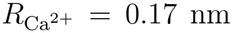, taken from Ref. [13], where these effective radii were validated by reproducing the radii of gyration of the *Azoarcus* ribozyme in Mg^2+^ and Ca^2+^. All-atom simulations of the *Azoarcus* ribozyme, in which Mg^2+^ retains its full hydration shell, yield the same Mg^2+^ binding sites as the coarse-grained model [13].

### Stacking Potential

Stacking between two adjacent nucleotides, *i* and *i*+1, is described by the potential [15, 17],

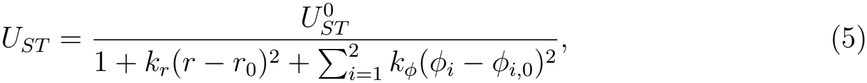

where *r* is the separation between the base sites B_i_ and B_i+1_. The angles *ϕ*_1_ (P_i_, S_i_, P_i+1_, S_i+1_) and *ϕ*_2_ (P_i+2_, S_i+1_, P_i+1_, S_i_) are the backbone dihedrals about the sugar–phosphate bonds. The reference values *r*_0_, *ϕ*_1,0_ and *ϕ*_2,0_ are obtained from the coarse-grained representation of an ideal B-form DNA helix. The parameter 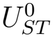 sets the strength of stacking between consecutive nucleotides in this ideal helix. At *T* = 277 K, its value is ≈ −4.19 kcal/mol for a T-dimer and −5.77 kcal/mol for an A-dimer (see Table S2) [15]. These values were calibrated previously [15] so that the model reproduces the thermodynamics of DNA duplexes predicted by the nearest-neighbor model [18, 19].

### Hydrogen bond interactions

*U*_*HB*_: Hydrogen bonding is included only between canonical Watson–Crick base pairs on complementary strands. The corresponding potential is

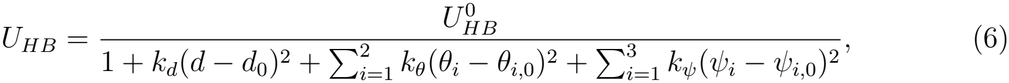

where the distance *d*, the angles *θ*_1_ and *θ*_2_, and the dihedral angles *ψ*_1_, *ψ*_2_ and *ψ*_3_ are defined in Figure 1E. The values of the parameters in *U*_*HB*_, and the procedure used to determine them, are given in our earlier work [15]. Because the ssDNA sequences studied here are homopolymers, they do not form hydrogen bonds.

### Electrostatic interactions

*U*_*EL*_: Electrostatic interactions between charged sites *i* and *j*, with charges *Q*_*i*_ and *Q*_*j*_ and separation *r*_*ij*_, are described by the Coulomb potential,

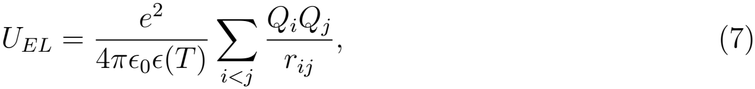

where *ϵ*_0_ is the permittivity of vacuum and *ϵ*(*T*) is the temperature-dependent dielectric constant of water [20],

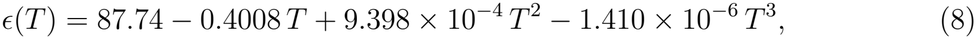

with *T* in degrees Celsius. Each phosphate group carries a charge *Q*_*P*_ = −1 in units of the elementary charge *e*.

### Model for

Spm^4+^: We modeled Spm^4+^ using seven coarse-grained interaction centers (Figure 1B). Four sites represent the amine groups, and the remaining three are the alkyl segments. The energy of a PA molecule is, *U*_*PA*_ = *U*_*B*_ + *U*_*EV*_ + *U*_*El*_. The bond stretch term, *U*_*B*_, is taken to be a harmonic potential, *U*_*B*_ = *k*_*b*_(*r* − *r*_0_)^2^. The equilibrium bond length, *r*_0_, is the sum of the radii of the bonded beads *i* and *j*, and *k*_*b*_ = 20.0 kcal/mol/Å^2^ is the spring constant. Excluded volume repulsion, *U*_*EV*_ , between any two non-bonded sites *i* and *j*, separated by a distance, *r* (in nm), is,

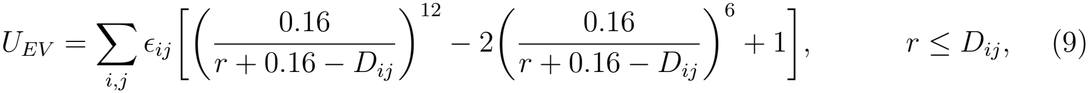

where *D*_*ij*_ = *R*_*i*_+*R*_*j*_ and 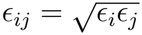 *R*_*i*_ and *R*_*j*_ are the radii of *i*^*th*^ and *j*^*th*^ beads, respectively, and *ϵ*_*i*_ and *ϵ*_*j*_ are the respective interaction strengths. The radius (=0.163 nm) and *ϵ* (=0.17 kcal/mol) values for the amine beads are taken from the OPLS-AA [21] force field parameters of amine nitrogen. We used the same *ϵ* values for the amine and the alkyl groups because the short-range nature of the repulsive excluded volume interaction should be insensitive to the *ϵ* values. The radii for the alkyl groups, (CH_2_)_3_ (=0.293 nm) and (CH_2_)_4_ (=0.308 nm), are taken from the Self-Organized Polymer model with side chains (SOP-SC) [22, 23], corresponding to the side chain radii of valine and leucine, respectively. The interactions between DNA and Spm^4+^ are governed by electrostatics and volume exclusion.

### Simulations

We used OpenMM [24] to simulate ssDNA on a Graphics Processing Unit (GPU). Long-range Coulomb interactions were treated using Particle-Mesh-Ewald (PME) algorithm [25]. In order to enhance conformational sampling, the simulations were performed by integrating the Langevin equation in the low friction limit [26]. The equation of motion for a bead or ion, with coordinate 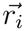, is,

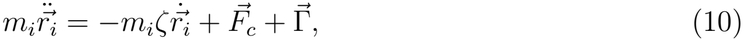

where *m*_*i*_ is the mass of the bead, *ζ* is the friction coefficient, 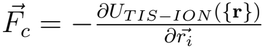, and 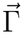 is the random force with a white noise spectrum. The autocorrelation function of the random force in the discretized form is 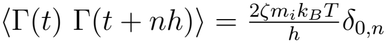, where *n* = 0, 1, … and *δ*_0,*n*_ is the Kronecker delta function. The equations of motion were integrated using Langevin leap-frog method [27] using *ζ* = 0.01 picosecond^−1^. The integration time step is 2 femtoseconds. We performed simulations at *T* = 277 K to mimic the ion counting experiments performed on the same constructs [28].

A single ssDNA polymer is placed in a cubic box of length *L* whose value depends on the ion concentration (*L* varied between 23 to 83 nm). To minimize finite size effects, we used periodic boundary conditions. Neutrality of the sample is maintained by adding an appropriate number of Cl^−^ ions. In all the simulations, 20 mM NaCl was also included.

## RESULTS

### Calculation of *l*_*p*_

We obtained the persistence length, *l*_*p*_, by fitting the simulated distribution of end-to-end distances (see Figure S1) to the distribution function for a semiflexible chain [29–31]. A mean-field expression for *P*(*r*_*ee*_, *L*) is,

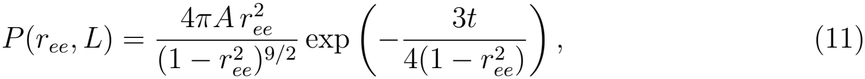

where *r*_*ee*_ = *R*_*ee*_*/L* is the end-to-end distance, *R*_*ee*_, normalized by the contour length, *L*, and *t* = *L/l*_*p*_. The normalization constant is

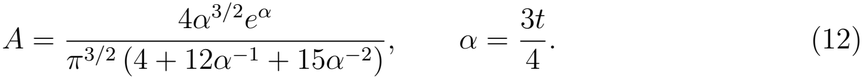

Murphy et al. used Eq. 13, together with FRET measurements, to estimate *l*_*p*_ of ssDNA in monovalent salt solutions [32]. We also computed *l*_*p*_ using an exact expression for *P*(*r*_*ee*_, *L*) [29] (see Supplemental Material for the comparison). As an additional check, we calculated *l*_*p*_ directly from the simulated 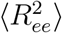 using the worm-like chain relation,

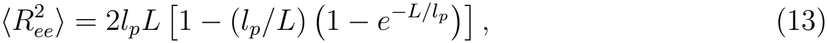

which does not depend on the shape of the distribution. Despite the poorer fits for dA_30_, the two estimates agree to within 5% for both sequences, and they yield the same dependence on ion concentration (Figure S2). Thus, the values of *l*_*p*_ reported here are robust to the method of estimation.

### Persistence length in divalent cations

The persistence length *l*_*p*_ of ssDNA decreases monotonically as the Mg^2+^ (or Ca^2+^) concentration increases (Figure 2). To test the applicability of PE theories, we fit the simulated *l*_*p*_ to Eq. 2. The fits (solid red lines in Figure 2) reveal a few notable trends. First, for dT_30_, the linear dependence of *l*_*p*_ on *κ*^−1^ holds in the presence of divalent cations, which is noteworthy because the assumptions (PE is flexible and the electrostatic interactions are described by the DH potential) leading to Eq. 2 [3, 4] are not usually satisfied in divalent cations. Second, the values of 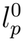 extracted from fits reflect the sequence-dependent intrinsic stiffness. The flexible dT_30_ 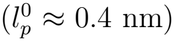 adopts a random-coil-like conformation (Figure 1D) due to weak stacking between adjacent bases. In contrast, dA_30_, which is more helical (Figure 1D) as a result of stronger base stacking, has a larger 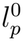 (≈ 1.1 nm). (3) The values of *l*_*p*_ are less in divalent cations than in monovalent salts [32], reflection the stronger interactions with the phosphate groups.

**FIG. 2:**
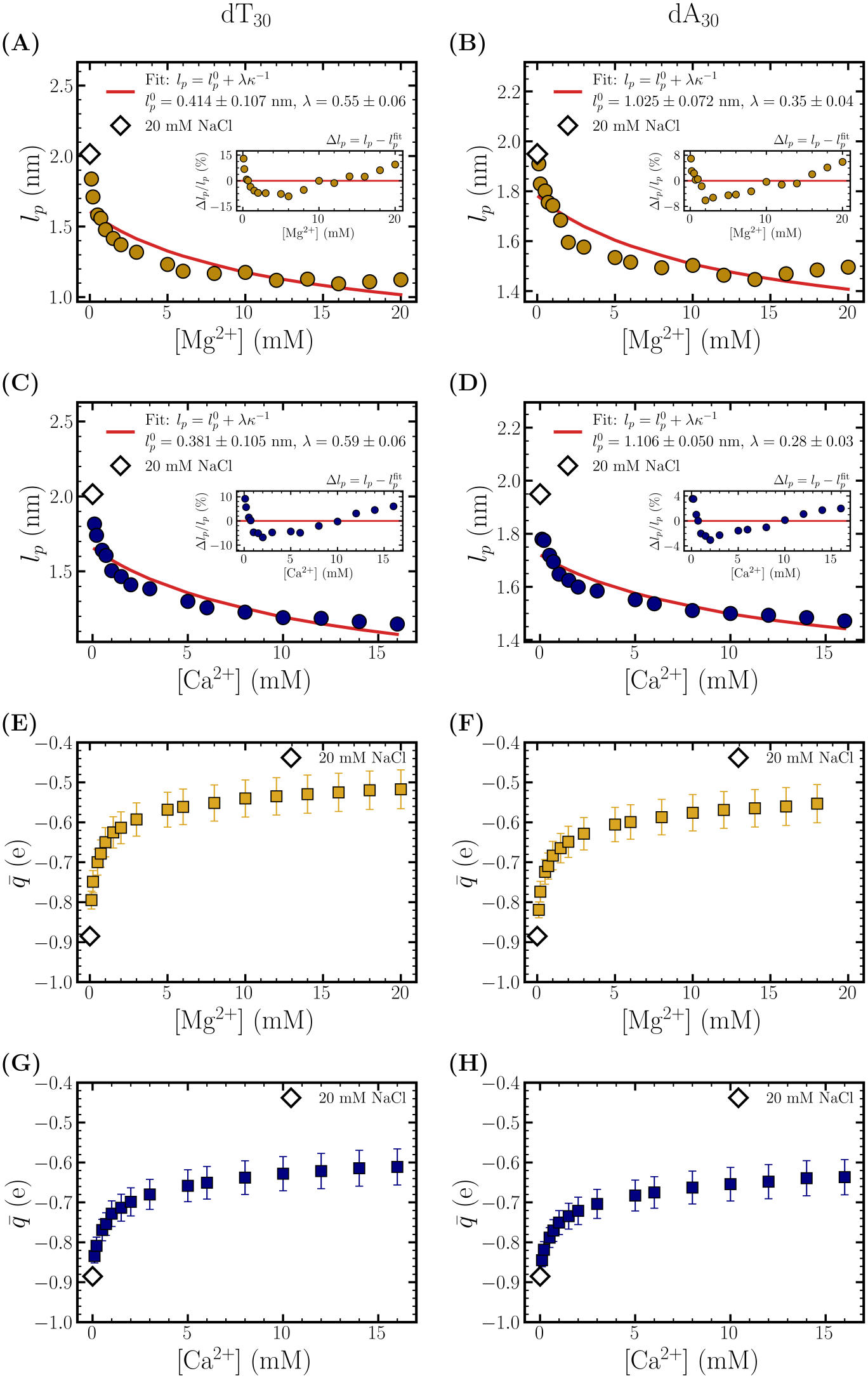
Persistence length in divalent cations: (A-B) Dependence of *l*_*p*_ as a function of Mg^2+^ concentration, in 20 mM NaCl. (A) is for dT_30_ and (B) is for dA_30_ (C, D) Same as (A, B), except the results are shown in Ca^2+^. The solid red curves are fits to the equation (Eq. 2). The parameters are specified in the legends. The insets show the relative deviation of the simulated *l*_*p*_ from the fit, Δ*l*_*p*_*/l*_*p*_, where 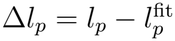. (E) Renormalized charge (see Eq. 14) on *P* groups of dT_30_ and (F) dA_30_, in Mg^2+^. (G, H) Same as (E, F), except the results are shown for Ca^2+^. 14

To distinguish between the *κ*^−1^ dependence (Eq. 2) and the *κ*^−2^ dependence predicted by the OSF theory (Eq. 1), we fixed *A* at the phosphate–phosphate distance (*A* = 0.65 nm, see Figure S3), so that each expression contains a single fitting parameter, 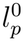 (Figure S4). The OSF expression fails for both sequences in Mg^2+^ and Ca^2+^. The residual, the sum of squares, is 5–14 times larger than predicted by Eq. 2. If the exponent of *κ*^−1^ is treated as a free parameter, we obtain *θ* = 0.89 *±* 0.08 (Mg^2+^) and 0.93 *±* 0.07 (Ca^2+^) for dT_30_ (Figure S5), which is in reasonable agreement with with Eq. 2. These results are in accord with the regimes of validity of the two theories [11]. The *κ*^−1^ dependence is expected when the OSF electrostatic persistence length exceeds the bare persistence length, whereas the OSF result is recovered in the opposite limit. For dT_30_, *l*_*OSF*_ ≈ 0.50–2.0 nm exceeds 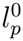 at all concentrations. In contrast, 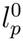 of dA_30_ is comparable to *l*_*OSF*_ . Thus, dA_30_ maybe in the crossover regime between the flexible (*κ*^−1^) and stiff (*κ*^−2^) limits. In accord with this expectation, the values of the fitted exponent for dA_30_ is *θ* ≈ 0.5 (Figure S5). We note that neither form captures the sharp decrease of *l*_*p*_ at low divalent ion concentrations, which is associated with the condensation of divalent ions onto the phosphate groups (Figure 2E–H).

A note on relation to experiments is worth making. The 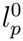 values obtained by fitting the simulated *l_p_* are comparable to the experimentally reported 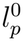 values (0.55 nm to 1.5 nm) for different ssDNA constructs, mostly in monovalent salts [5, 32–41]. Interestingly, *l*_*p*_ extracted from force extension traces generated in single molecule pulling experiments [39] on a long (6770 nucleotides) ssDNA sequence for a limited number of Mg^2+^ data points was fit using Eq. 2 with 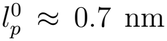. We hope that our work encourages additional experiments on ssDNA homopolymers in Mg^2+^ and Ca^2+^.

### Charge renormalization

Although *l*_*p*_ decreases as *κ*^−1^ (Eq. 2), the reduction is not uniform over the concentration range. In 20 mM NaCl, without divalent cations, *l*_*p*_ ≈ 2.0 nm for both sequences. The decrease in *l*_*p*_ is substantial upon addition of Mg^2+^ and Ca^2+^ up to ≈ 1 mM. This is followed by a moderate change in *l*_*p*_ at higher concentrations. The nonuniform decrease could be qualitatively explained using the counterion condensation theory [42, 43].

We used simulations to calculate the number of monovalent and divalent ions that are condensed onto the phosphate backbone. This enables the calculation of the renormalized charge 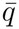 per phosphate group. We assume that a counterion is condensed onto a phosphate if the ion is within a cutoff distance *r*_*c*_ = (*r*_*P*_ + *r*_*i*_ + Δ*r*), where *r*_*i*_ is the radius of the *i*th bead and Δ*r* is taken to be 0.15 nm. The renormalized charge is computed using,

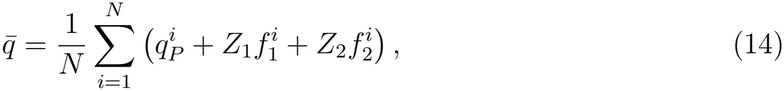

where 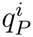 is the bare charge on the *i*^*th*^ phosphate group (−1), 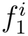 is the number of monovalent and 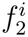 is the number of divalent ions bound to the *i*th phosphate, and *Z*_1_ and *Z*_2_ are the valencies of the monovalent and divalent ions, respectively.

In 20 mM NaCl solution, with no added Mg^2+^ ions, 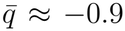 for both the sequences. Addition of Mg^2+^ up to 1 mM accounts for ≈ 25% charge reduction, while further addition of ions up to 20 mM leads to only ≈ 13% charge reduction. Thus, the two regimes in *l*_*p*_ mirror those in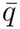. The value of 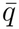 saturates at ≈ 50% charge reduction. In agreement with counterion condensation theory, the majority of the charge neutralization is due to condensed Mg^2+^ ions (≈ 47% for dT_30_ at the highest concentration), while Na^+^ neutralizes only ≈ 1% of the phosphate charges.

Following Oosawa-Manning theory [42, 44], we computed the renormalized charge on the phosphate using *q*_*OM*_ = *b/l*_*B*_(*T*), where *l*_*B*_(*T*) is the Bjerrum length and *b* is the length per unit charge, taken as 0.44 nm for single-stranded polynucleotides [45]. We estimate that *q*_*OM*_ is −0.6*e*, while from our simulations we obtain values close to −0.5*e* for dT_30_ and −0.6*e* for dA_30_. It follows that Mg^2+^ is more efficient in neutralizing the charge on dT_30_ than on dA_30_.

### Persistence length in spermine

We performed simulations to estimate the dependence of *l*_*p*_ on spermine concentration, which was varied from 0.1 mM to 20 mM. Interestingly, *l*_*p*_ does not change beyond 0.5 mM (Figure 3) in both dT_30_ and dA_30_. The lack of change in *l*_*p*_ even when the Spm^4+^ concentration is increased by more than two orders of magnitude (0.1 mM to 20 mM) is striking. Most of the decrease in *l*_*p*_ occurs at lower concentrations. For dT_30_, *l*_*p*_ decreases from 2.02 nm in 20 mM NaCl alone to 1.38 nm at 0.1 mM Spm^4+^, and then only to 1.15–1.18 nm for [Spm] ≥ 0.5 mM. Thus, ≈ 75% of the total decrease takes place below 0.1 mM. For dA_30_, the corresponding values are 1.95 nm, 1.65 nm and 1.52–1.59 nm. In contrast, 0.1 mM Mg^2+^ accounts for only ≈ 20% (dT_30_) and ≈ 10% (dA_30_) of the total decrease. At all concentrations, *l*_*p*_ is larger for dA_30_ than for dT_30_. In the absence of theory to account for the anisotropic nature of spermine molecule with four charges, there is no clear explanation of the simulation results.

**FIG. 3:**
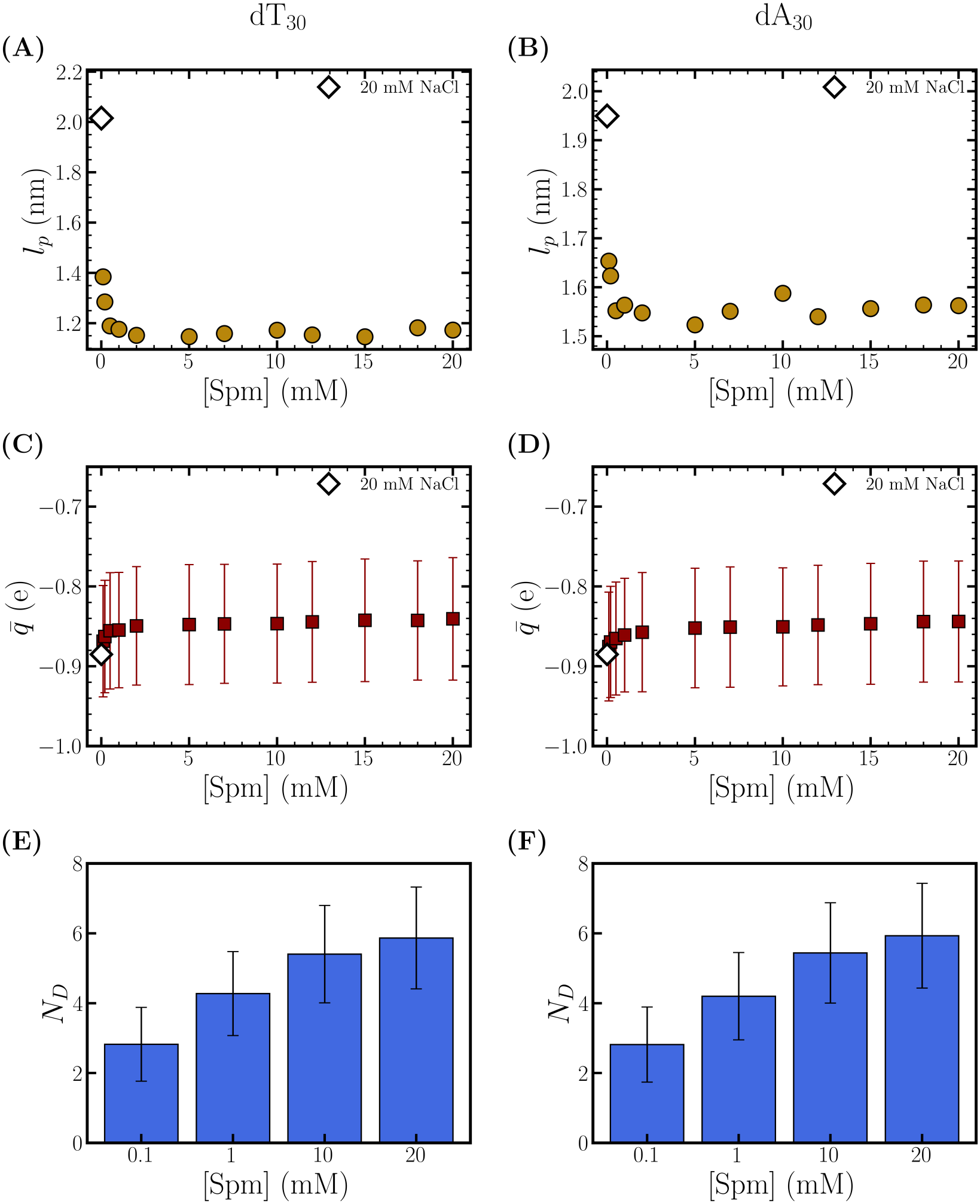
Persistence length in spermine: (A) Dependence of *l*_*p*_ of dT_30_ and (B) dA_30_ as a function of Spm concentration in 20mM NaCl. (C) Renormalized charge (see Eq. 14) on *P* groups of dT_30_ and (D) dA_30_. (E) Number of Spm molecules (*N*_*D*_) in direct contact with at least one of the phosphate (*P*) groups, for dT_30_ and (F) dA_30_. A direct contact means at least one of the amine moiety in Spm and the *P* groups are within the Bjerrum length.

The lack of change in *l*_*p*_ is also reflected in the renormalized charge 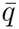 (Eq. 14). It also saturates near [Spm] ≈ 0.5 mM (Figures 3C,D). At all concentrations, 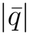 remains larger than in Mg^2+^ and Ca^2+^, indicating weaker charge neutralization by Spm. Nevertheless, at the highest concentrations, *l*_*p*_ in Spm^4+^ is nearly the same as in Mg^2+^ and Ca^2+^ (≈ 1.1–1.2 nm for dT_30_ and ≈ 1.5–1.6 nm for dA_30_), and the numbers of bound cations are also comparable. Thus, *l*_*p*_ at high concentration is not determined by charge neutralization alone.

These trends point to important differences between spermine and divalent cations. At neutral pH, spermine has four protonated amine groups linked along a flexible back-bone, unlike spherical divalent cations such as Mg^2+^ and Ca^2+^. This connectivity produces anisotropic interactions with the nucleotides and enhances the effects of excluded volume. A bound Spm^4+^ could hinder binding to the neighboring phosphate sites, limiting the number of spermine molecules that bind to ssDNA simultaneously. To illustrate the importance of spermine volume exclusion, we define a direct Spm–DNA contact when at least one amine group is within *l*_*B*_ (≈ 0.7 nm) of a phosphate group. Let *N*_*D*_ be the total number of such bound molecules. Figures 3E and F show that *N*_*D*_ increases only gradually, from ≈ 2.8 to ≈ 5.9, over a 200-fold increase in spermine concentration (0.1–20 mM).

We fit *N*_*D*_ to the Langmuir isotherm[46] and to the McGhee–von Hippel model[47], in which each bound Spm occupies *n* consecutive nucleotides (Figure S8). The McGhee–von Hippel fits give *n* = 5 for both sequences. Thus, each bound Spm excludes other Spm molecules from about five nucleotides, which limits binding to ≈ 6 molecules per chain. However, neither model captures the concentration dependence of *N*_*D*_. Both predict saturation near 1 mM, whereas *N*_*D*_ continues to increase up to 20 mM. The models assume that the affinity of Spm is independent of the number of molecules already bound. Each bound Spm^4+^, however, reduces the effective charge on the chain. As a result, the affinity for subsequent Spm molecules could account for the gradual increase in *N*_*D*_. Because most of the changes in *l*_*p*_ occurs once the first few Spm molecules bind, the additional binding at higher concentrations has little effect on the ssDNA conformations, hence on *l*_*p*_.

Figure 4 shows representative all-atom conformations of dT_30_ and dA_30_ with bound Spm molecules in 0.1 mM and 20 mM Spm. Despite the 200-fold increase in Spm concentration, the number of bound Spm molecules increases only modestly, which is consistent with the results in Figures 3E and F. We generated the all-atom structures by mapping the coarse-grained conformations to the atomically detailed representation [16].

**FIG. 4:**
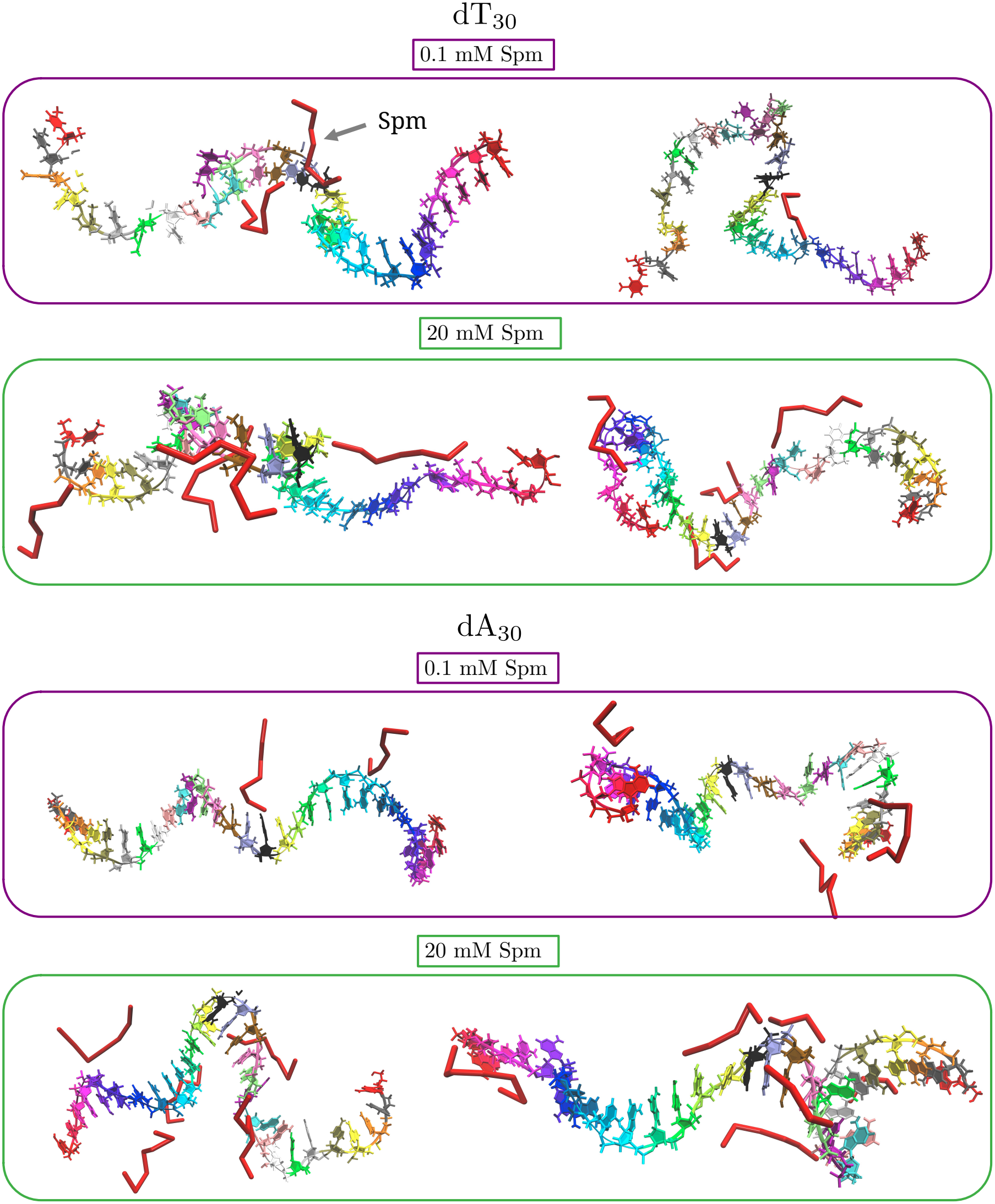
ssDNA conformations in spermine: Reconstructed all-atom structures for dT_30_ and dA_30_, obtained from coarse-grained simulations in 0.1 mM and 20 mM Spm. To aid visualization, nucleotides are assigned different colors based on the sequence number. Bound spermine molecules are shown in red.

To our knowledge, there are no experiments on the persistence length as a function of spermine concentration. Single-molecule force spectroscopy or FRET-based end-to-end distance measurements, similar to those used for divalent cations, could test our predictions. Such experiments would also elucidate whether excluded volume limits the binding of polyamines to ssDNA in general.

## CONCLUSIONS

We calculated the persistence length of short ssDNA chains (dT_30_ and dA_30_) in polyvalent cations using simulations of the coarse-grained TIS-DNA model with explicit ions[16]. In divalent cations, the electrostatic persistence length (*l*_*e*_) of dT_30_ is proportional to the Debye length, which unexpectedly agrees with theories developed for flexible polyelectrolytes. In contrast to dT_30_, neither the *κ*^−1^ nor the *κ*^−2^ dependence describes dA_30_ quantitatively. Because of strong base stacking, the bare persistence length of dA_30_ is comparable to the electrostatic persistence length, it is likely that neither form accounts for the changes in *l*_*p*_ as a function of cation concentration.

In contrast, we find that *l*_*p*_ is almost independent of Spm^4+^ concentration above 0.1 mM. There is no theoretical explanation for this unexpected prediction. It should be pointed out that binding of Spm^4+^ to the phosphate groups is favorable. However, it is accompanied by a loss in translational and rotational entropy of spermine. Importantly, the number of Spm^4+^ that can bind to ssDNA is restricted because of the large size of the cation. Binding increases only gradually with increasing concentration. Given the importance of polyamines in general and spermine in particular, experiments are needed to validate our predictions.

## Acknowledgments

This article is a tribute to M. Muthukumar whose numerous works in polymer physics have been a source of inspiration to DT. We are grateful to NSF (grant CHE 2320256) and the Welch Foundation (grant F-0019), administered through the Collie-Welch Regents Chair, for supporting this research. We thank the Texas Advanced Computing Center and Pittsburgh Supercomputing Center for providing generous computational resources.

## Data Availability

The data that supports the findings of this study are available from the corresponding author upon reasonable request.

