## Supplementary Figures S1-S8 and Tables S1-S2 for "Persistence length of short homopolymeric single-stranded DNA sequences in polyvalent cations"

### Supporting Information for "Persistence length of short homopolymeric single-stranded DNA sequences in polyvalent cations"

Balaka Mondal[†] and D. Thirumalai[*,†]

†*Department of Chemistry, The University of Texas at Austin, Austin, TX 78712, USA*

‡*Department of Physics, The University of Texas at Austin, Austin, TX 78712, USA*

¶*Department of Chemistry, University of Texas, Austin, TX, USA*

The SI we provide additional results to support of the results in the main text. They mostly deal with the goodness of fits of the end-to-end distribution functions, which was one of ways used to extract the persistence length $l_p$ as a function of cation concentration. In addition, we provide details of the fits of $l_p$ to cation concentration and theories developed for PE.

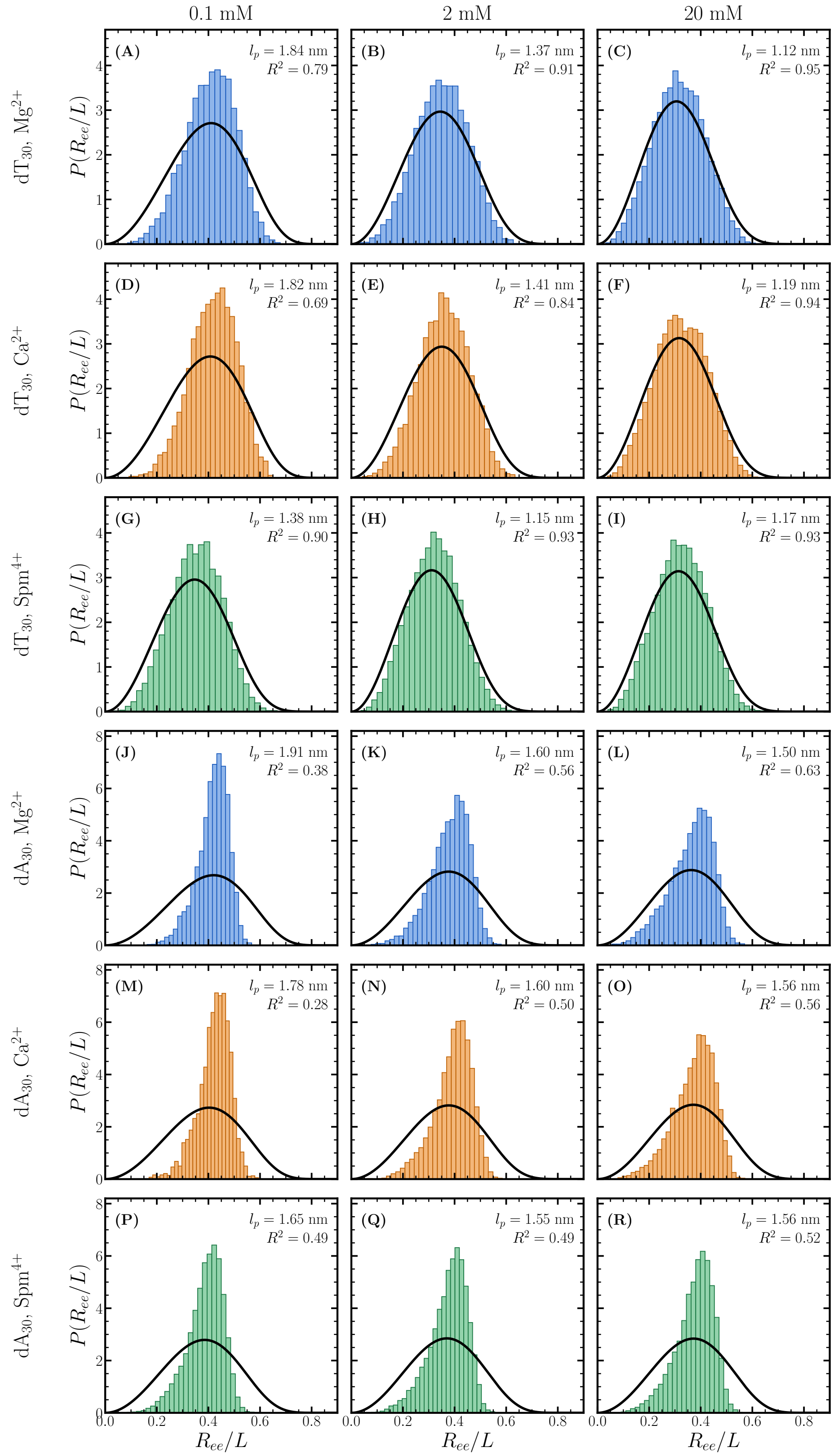
0.1 mM
2 mM
20 mM
dT30, Mg2+
dT30, Ca2+
dT30, Spm4+
dA30, Mg2+
dA30, Ca2+
dA30, Spm4+
P(Ree/L)
Ree/L
(A) lp = 1.84 nm R2 = 0.79
(B) lp = 1.37 nm R2 = 0.91
(C) lp = 1.12 nm R2 = 0.95
(D) lp = 1.82 nm R2 = 0.69
(E) lp = 1.41 nm R2 = 0.84
(F) lp = 1.19 nm R2 = 0.94
(G) lp = 1.38 nm R2 = 0.90
(H) lp = 1.15 nm R2 = 0.93
(I) lp = 1.17 nm R2 = 0.93
(J) lp = 1.91 nm R2 = 0.38
(K) lp = 1.60 nm R2 = 0.56
(L) lp = 1.50 nm R2 = 0.63
(M) lp = 1.78 nm R2 = 0.28
(N) lp = 1.60 nm R2 = 0.50
(O) lp = 1.56 nm R2 = 0.56
(P) lp = 1.65 nm R2 = 0.49
(Q) lp = 1.55 nm R2 = 0.49
(R) lp = 1.56 nm R2 = 0.52

Figure S1

Figure S1 caption: Distributions of $R_{ee}/L$ for $dT_{30}$ (A–I) and $dA_{30}$ (J–R) at 0.1, 2 and 20 mM $Mg^{2+}$, $Ca^{2+}$ and $Spm^{4+}$. Black lines are fit to the Eq. 11 in the main text. $l_p$ values are reported in each panel.

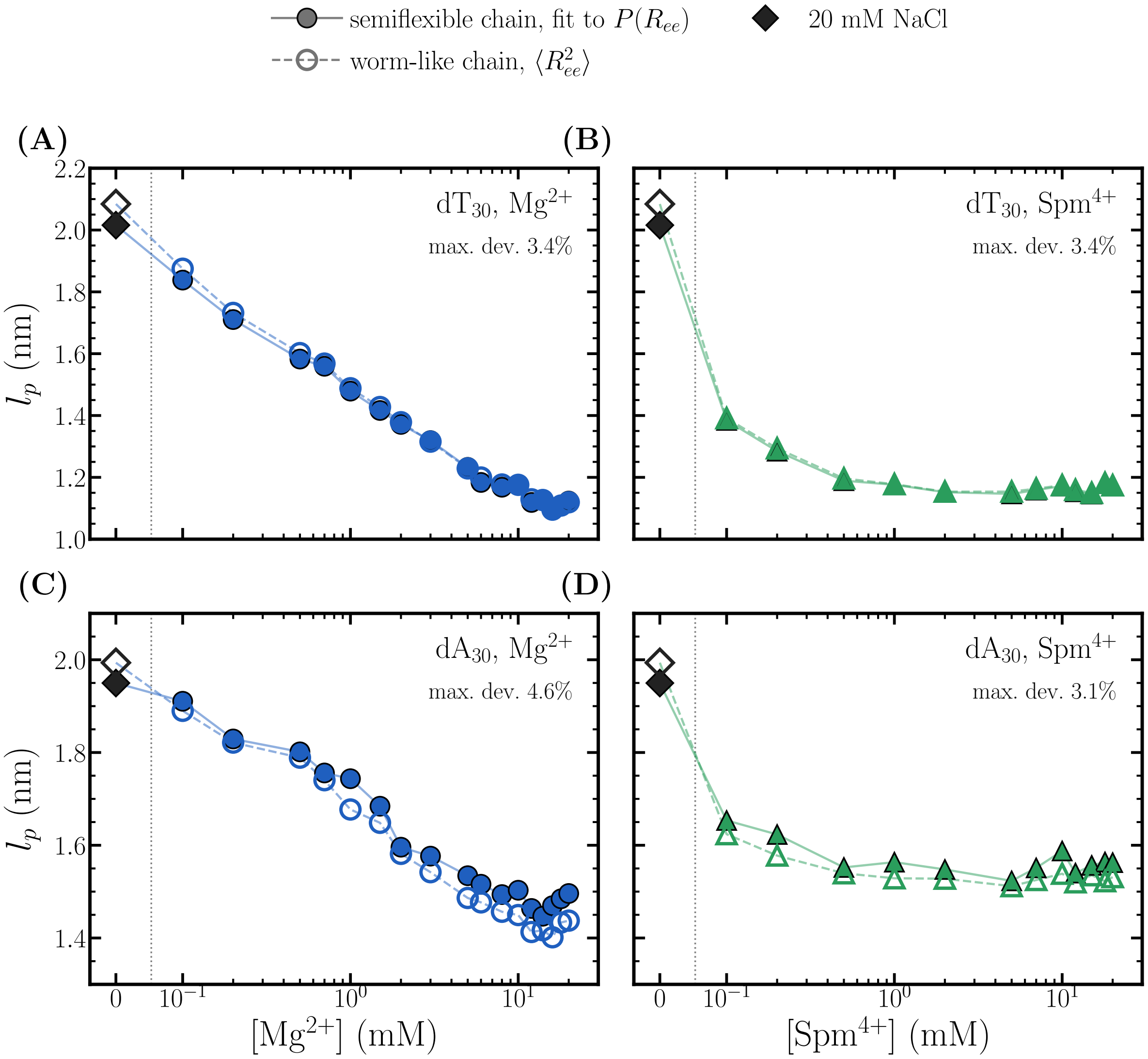


Figure S2: Persistence length of $dT_{30}$ (A, B) and $dA_{30}$ (C, D) as a function of $[Mg^{2+}]$ (A, C) and $[Spm^{4+}]$ (B, D). Filled symbols (solid lines): $l_p$ obtained by fitting the simulated distribution of the end-to-end distance, $P(R_{ee})$, to the semiflexible-chain model. Open symbols (dashed lines): $l_p$ obtained from the simulated mean-square end-to-end distance using the worm-like-chain relation $\langle R_{ee}^2 \rangle = 2l_pL\left[1 - \frac{l_p}{L}\left(1 - e^{-L/l_p}\right)\right]$.

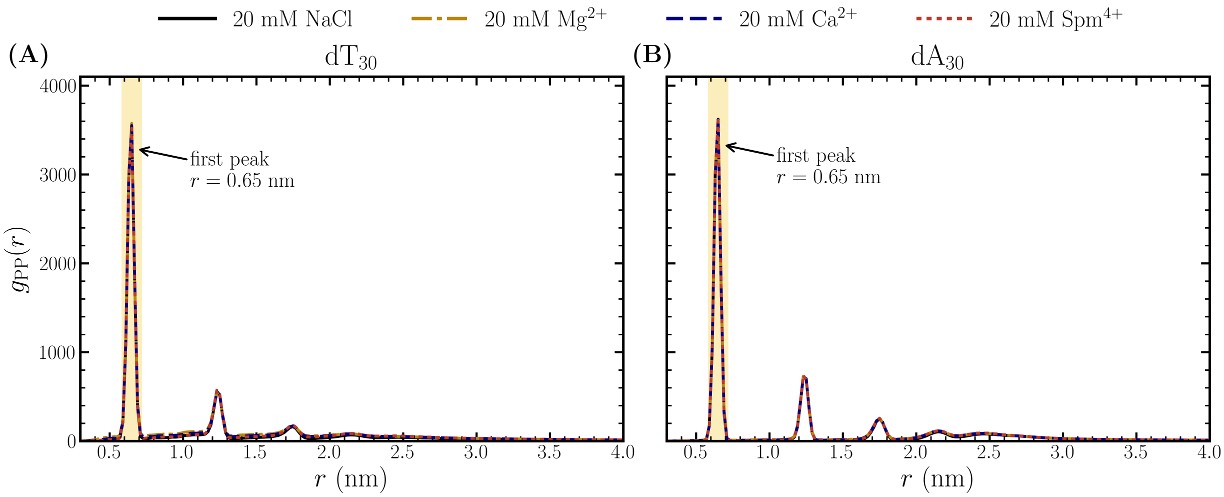


Figure S3: Radial distribution functions, $g_{PP}(r)$ between phosphate groups of $dT_{30}$ and $dA_{30}$ at different ion concentrations.

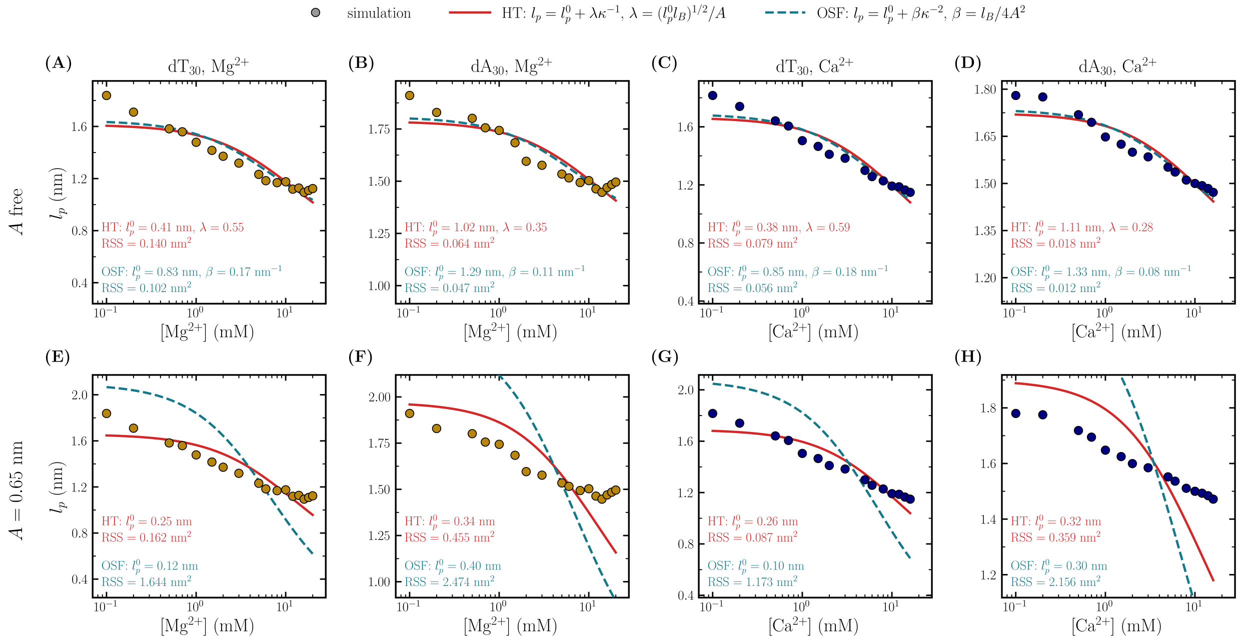


Figure S4: Persistence length of $dT_{30}$ and $dA_{30}$ as a function of $Mg^{2+}$ and $Ca^{2+}$ concentration (20 mM NaCl), fit to $l_p = l_p^0 + \lambda\kappa^{-1}$ (Eq. 2, solid) and $l_p = l_p^0 + \beta\kappa^{-2}$ (OSF, dashed), with $\lambda = (l_p^0 l_B/A^2)^{1/2}$, $\beta = l_B/4A^2$ and $l_B = 0.70$ nm. (A–D) $A$ treated as a free parameter ($l_p^0$ and $\lambda$ or $\beta$ fitted). (E–H) $A = 0.65$ nm, the phosphate–phosphate distance ($l_p^0$ fitted). The fit parameters and residual sums of squares (RSS) are given in each panel.

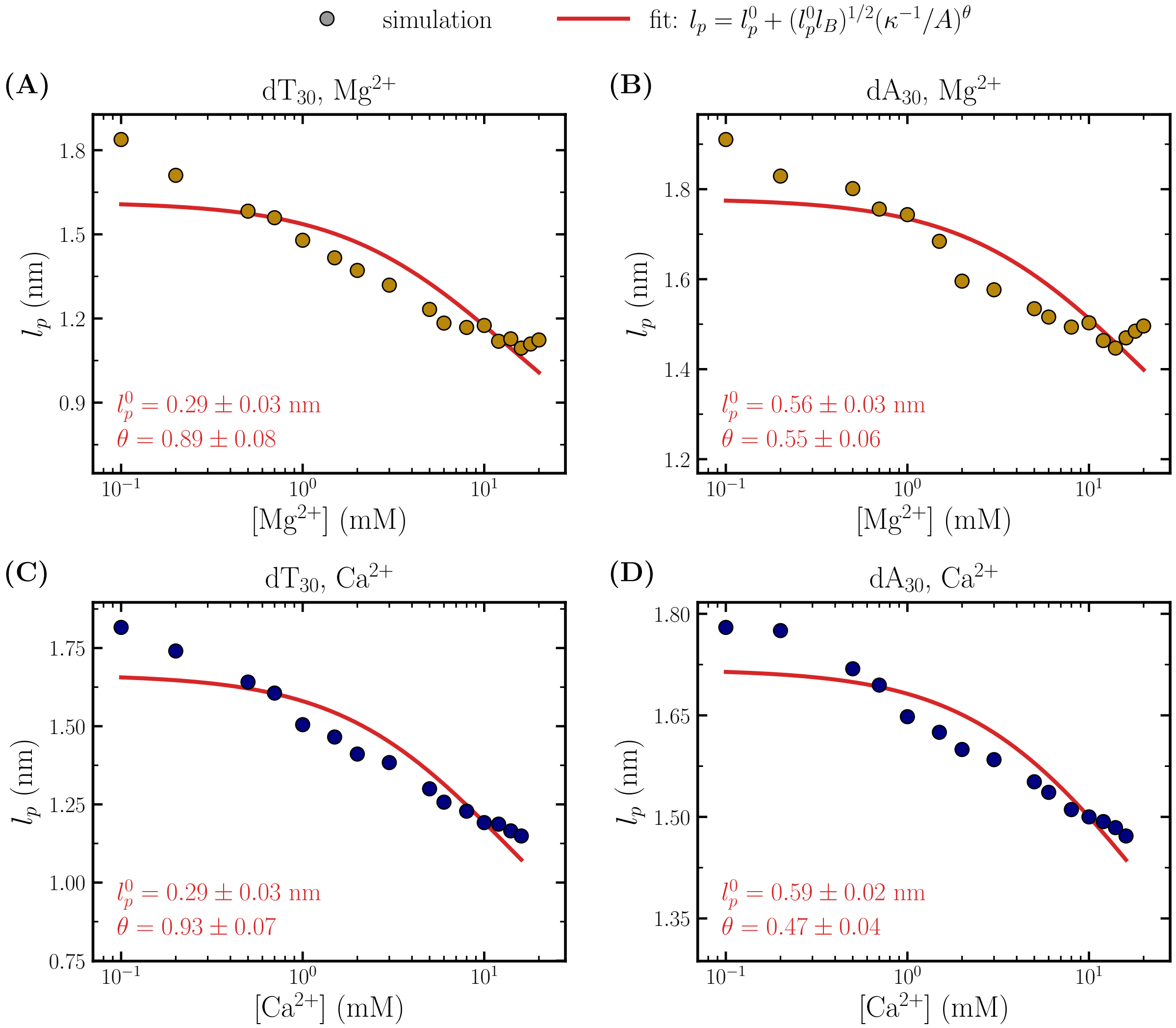


Figure S5: Persistence length of $dT_{30}$ and $dA_{30}$ in (A, B) $Mg^{2+}$ and (C, D) $Ca^{2+}$ (20 mM NaCl). Solid lines are fits to $l_p = l_p^0 + (l_p^0 l_B)^{1/2}(\kappa^{-1}/A)^\theta$ with $A = 0.65$ nm and $l_B = 0.70$ nm, and with $l_p^0$ and $\theta$ as fit parameters. For $\theta = 1$ the expression reduces to Eq. 2.

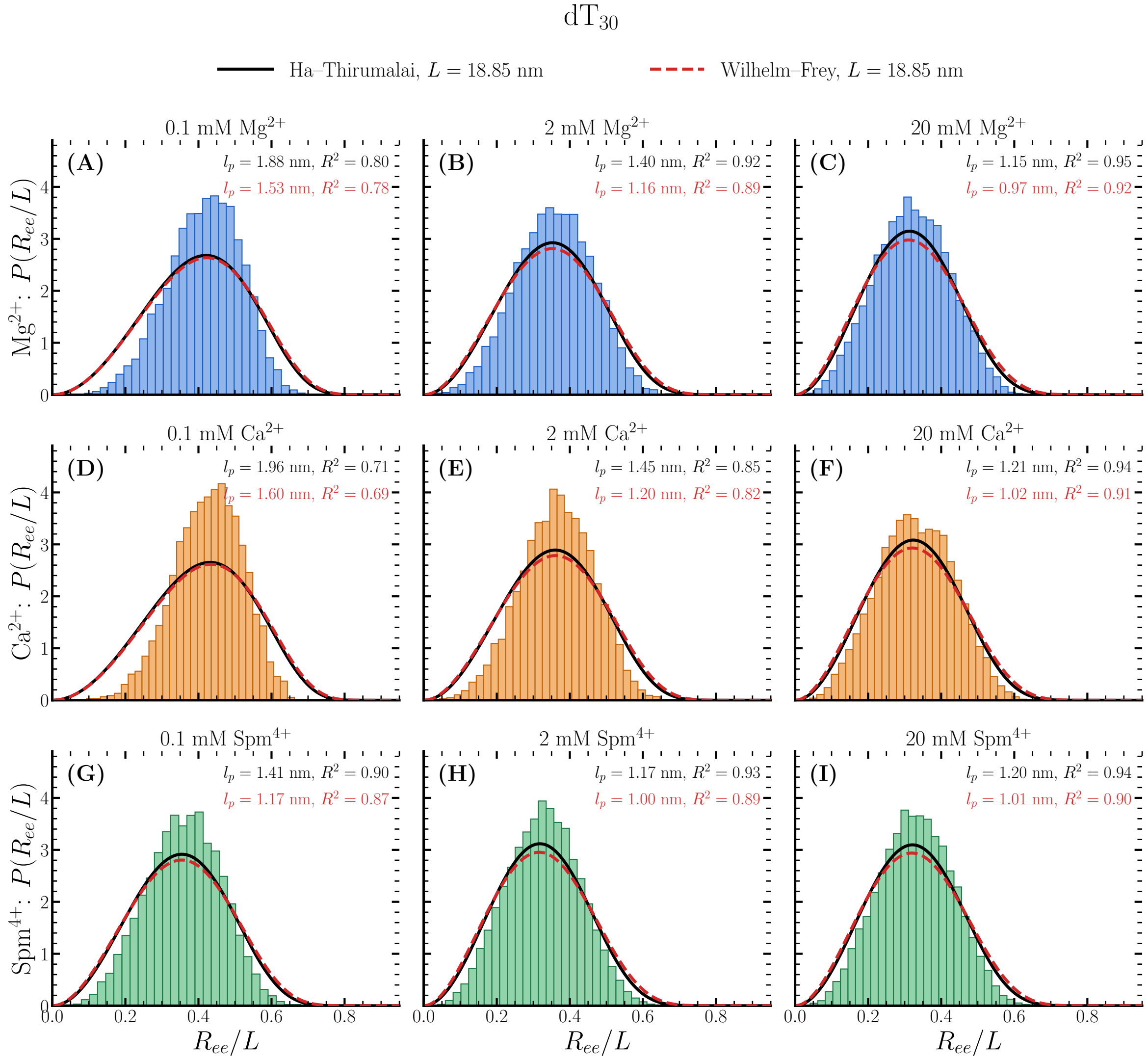


Figure S6: Distributions of the normalized end-to-end distance, $R_{ee}/L$, of $dT_{30}$ in (A–C) $Mg^{2+}$, (D–F) $Ca^{2+}$ and (G–I) $Spm^{4+}$ at 0.1, 2 and 20 mM, with 20 mM NaCl in all cases. Histograms are simulation data. Solid lines are fits to the Ha–Thirumalai distribution.[1,2] Dashed lines are fits to the exact result obtained by Wilhelm–Frey for semi-flexible chains.[3] The fits used both with same contour length $L$. The fitted persistence lengths, $l_p$, $R^2$, is given in each panel for both fits.

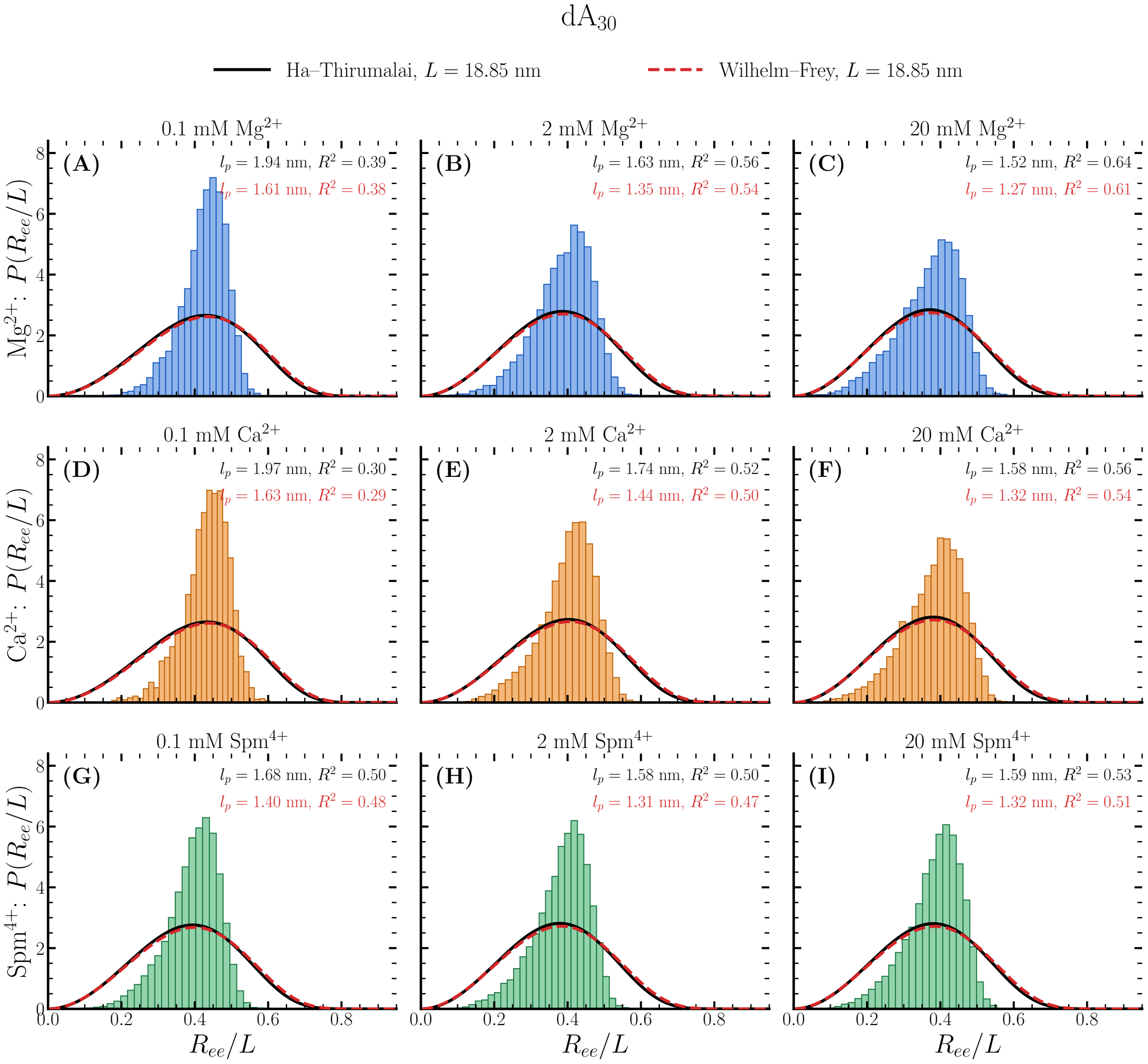


Figure S7: Same as Figure S6, except the results are for dA$_{30}$.

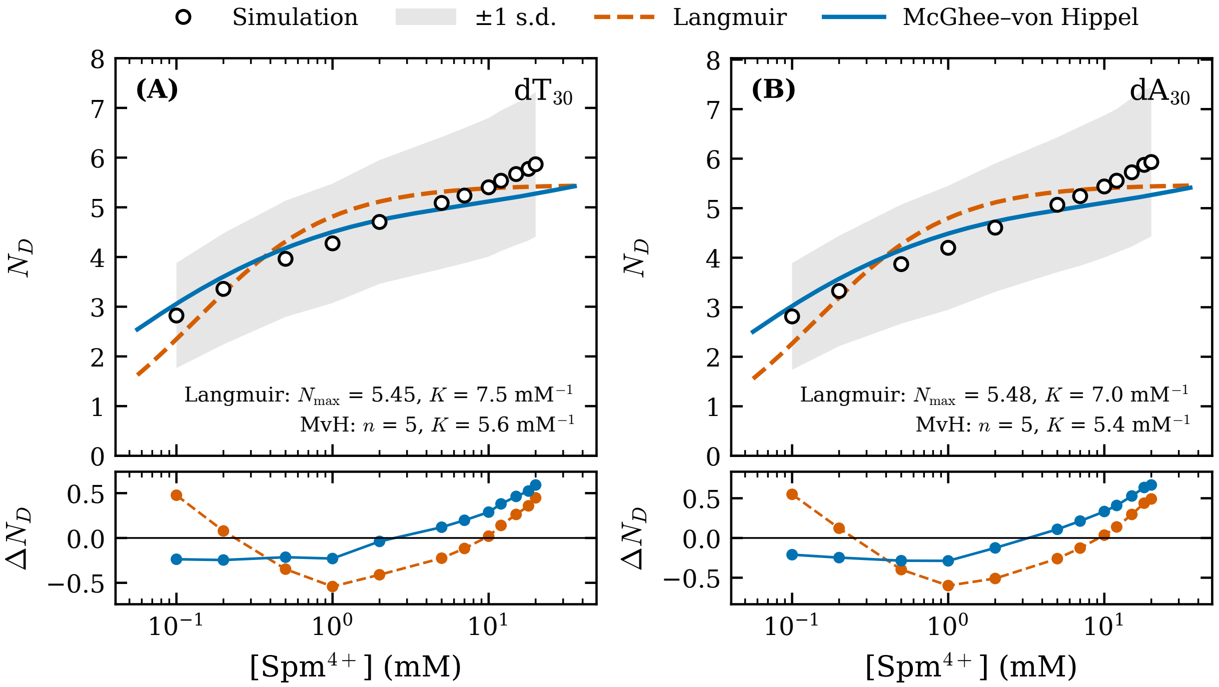


Figure S8: Number of Spm molecules ($N_D$) in direct contact with at least one of the phosphate ($P$) groups as a function of Spm concentration in 20 mM NaCl, for (A) $dT_{30}$ and (B) $dA_{30}$. Circles are simulation results, and the shaded region shows ±1 standard deviation of $N_D$. The dashed and solid lines are least-squares fits to the Langmuir isotherm, $N_D = N_{\max}Kc/(1 + Kc)$, and to the McGhee–von Hippel model for a chain of $N = 30$ nucleotides, in which each bound Spm occupies $n$ consecutive nucleotides. The fit parameters are given in each panel. The lower panels show the residuals, $\Delta N_D = N_D^{\rm sim} - N_D^{\rm fit}$.

**Table S1: Parameters for excluded volume and electrostatic interactions**

| | $R_i$ nm | $\epsilon_i$ kcal/mol | $q_i$ e |
|---|---|---|---|
| P | 0.21 | 0.2 | -1 |
| S | 0.29 | 0.2 | 0 |
| A | 0.28 | 0.2 | 0 |
| G | 0.3 | 0.2 | 0 |
| T | 0.27 | 0.2 | 0 |
| C | 0.27 | 0.2 | 0 |
| $Mg^{2+}$ | 0.08 | 0.9 | 2 |
| $Ca^{2+}$ | 0.17 | 0.5 | 2 |
| $Na^{+}$ | 0.19 | 0.00277 | 1 |
| $Cl^{-}$ | 0.19 | 0.3 | -1 |

**Table S2: Parameters to compute neighboring stacking energy (Eq. 5 of main text), $U^0_{ST} = -h + k_{\mathrm{B}}(T - T_m)s$.**

| $\frac{x}{w}$ | $h$ (kcal/mol) | $s$ | $T_m$ (K) |
|---|---|---|---|
| $\frac{A}{A}$ | 5.69 | 0.94 | 322.0 |
| $\frac{A}{T}$; $\frac{T}{A}$ | 4.95; 5.03 | 0.87; 0.65 | 293.0 |
| $\frac{A}{G}$; $\frac{G}{A}$ | 5.43; 5.42 | 1.06; 0.98 | 333.6 |
| $\frac{A}{C}$; $\frac{C}{A}$ | 4.98; 5.03 | 0.92; 0.79 | 293.0 |
| $\frac{G}{G}$ | 5.66 | −0.29 | 353.9 |
| $\frac{G}{C}$; $\frac{C}{G}$ | 5.22; 5.13 | 2.33; 2.41 | 331.9 |
| $\frac{G}{T}$; $\frac{T}{G}$ | 5.28; 5.13 | 1.69; 1.73 | 332.6 |
| $\frac{C}{C}$ | 4.15 | 0.98 | 288.3 |
| $\frac{C}{T}$; $\frac{T}{C}$ | 4.13; 4.18 | 0.71; 0.94 | 288.3 |
| $\frac{T}{T}$ | 4.17 | 0.89 | 288.3 |

$k_d = 1.45$ Å$^{-2}$, $k_\psi = 3.0$ rad$^{-2}$;
